# SUsPECT: A pipeline for variant effect prediction based on custom long-read transcriptomes for improved clinical variant annotation

**DOI:** 10.1101/2022.10.23.513417

**Authors:** Renee Salz, Nuno Saraiva-Agostinho, Emil Vorsteveld, Caspar I. van der Made, Simone Kersten, Merel Stemerdink, Jamie Allen, Pieter-Jan Volders, Sarah E. Hunt, Alexander Hoischen, Peter A.C. ’t Hoen

## Abstract

Our incomplete knowledge of the human transcriptome impairs the detection of disease-causing variants, in particular in transcripts only expressed under certain conditions. These transcripts are often lacking from reference transcript sets, such as Ensembl/GENCODE and RefSeq, and could be relevant for establishing genetic diagnoses. We present SUsPECT (Solving Unsolved Patient Exomes/gEnomes using Custom Transcriptomes), a pipeline based on the Ensembl Variant Effect Predictor (VEP) to predict variant impact on custom transcript sets, such as those generated by long-read RNA-sequencing, for downstream prioritization. Our pipeline predicts the functional consequence and likely deleteriousness scores for missense variants in the context of novel open reading frames predicted from any transcriptome. We demonstrate the utility of SUsPECT by uncovering potential mutational mechanisms of pathogenic variants in ClinVar that are predicted to be benign using the reference transcript annotation. In further support of SUsPECT’s utility, we identified an enrichment of immune-related variants predicted to have a more severe molecular consequence when annotating with a newly generated transcriptome from stimulated immune cells instead of the reference transcriptome. Our pipeline outputs crucial information for further prioritization of potentially disease-causing variants for any disease and will become increasingly useful as more long-read RNA sequencing datasets become available.

## Background/objectives

The number of recorded nucleic acid variants in the human genome increased significantly with the advent of next-generation sequencing (NGS). The sequencing of the new genetic variants has outpaced the understanding of them. As genetic diversity is linked to disease susceptibility, therapy response and clinical outcomes, there is great interest in accurately predicting the functional consequences of genetic variants. Since only a small fraction of all available variants can be characterized clinically or by functional efforts, there is a heavy reliance on computational methodology for prioritization. Several computational methods predict the effect of genetic variant effects on function such as PolyPhen-2 (1), SIFT (2), and MutPred2 (3). Variant annotators such as the Ensembl Variant Effect Predictor (VEP) (4) and ANNOVAR (5) collect gene/transcript information from reference databases (containing pre-computed scores of the aforementioned software in some cases) and provide effect predictions to end users. Their interpretation of the variant effects has implications for clinical diagnosis and treatment, and paves the way for precision medicine.

Short-read RNA sequencing has provided us with the majority of knowledge we currently have about the transcriptome, but has some intrinsic limitations with isoform discovery (6). As a result, the use of current reference transcript sequences does not provide a complete picture of how a variant affects molecular functioning. Long-read sequencing allows for the accurate elucidation of isoforms (7) and long-read RNA sequencing datasets are proving that the human transcriptome has much more diversity than previously thought (8–10). In addition, both short and long-read sequencing have shown that gene expression is highly variable in a context dependent manner, e.g. based on conditions (infection, stress, disease) or tissue- or cell-types (11–14).

Understanding the coding potential of these newly discovered transcripts is key to predicting functional consequences of variants within them. Since long-reads often capture whole transcripts, more accurate open reading frames (ORFs) can be predicted. Alternative splicing is known to increase the proteomic diversity, but less understood is the contribution of novel transcripts to this diversity and what it means for function (15–18). There are several computational methods available to predict ORFs of these novel transcripts either based on sequence features (19–21) or homology to existing protein coding transcripts (22–24). The prediction of ORFs on novel sequences is an essential first step for the detection of new proteoforms, as proteomics usually relies on previously observed sequences. Transcripts derived from long-read sequencing can provide better predictions of (novel) proteoforms (Figure 1). Thus, long-read transcriptome data relevant to the disease of interest may not only improve our understanding of the ever-growing number of genetic variants that are identified in human disease context, but also aid in diagnoses for rare and/or unsolved disease (25, 26).

**Figure 1:**
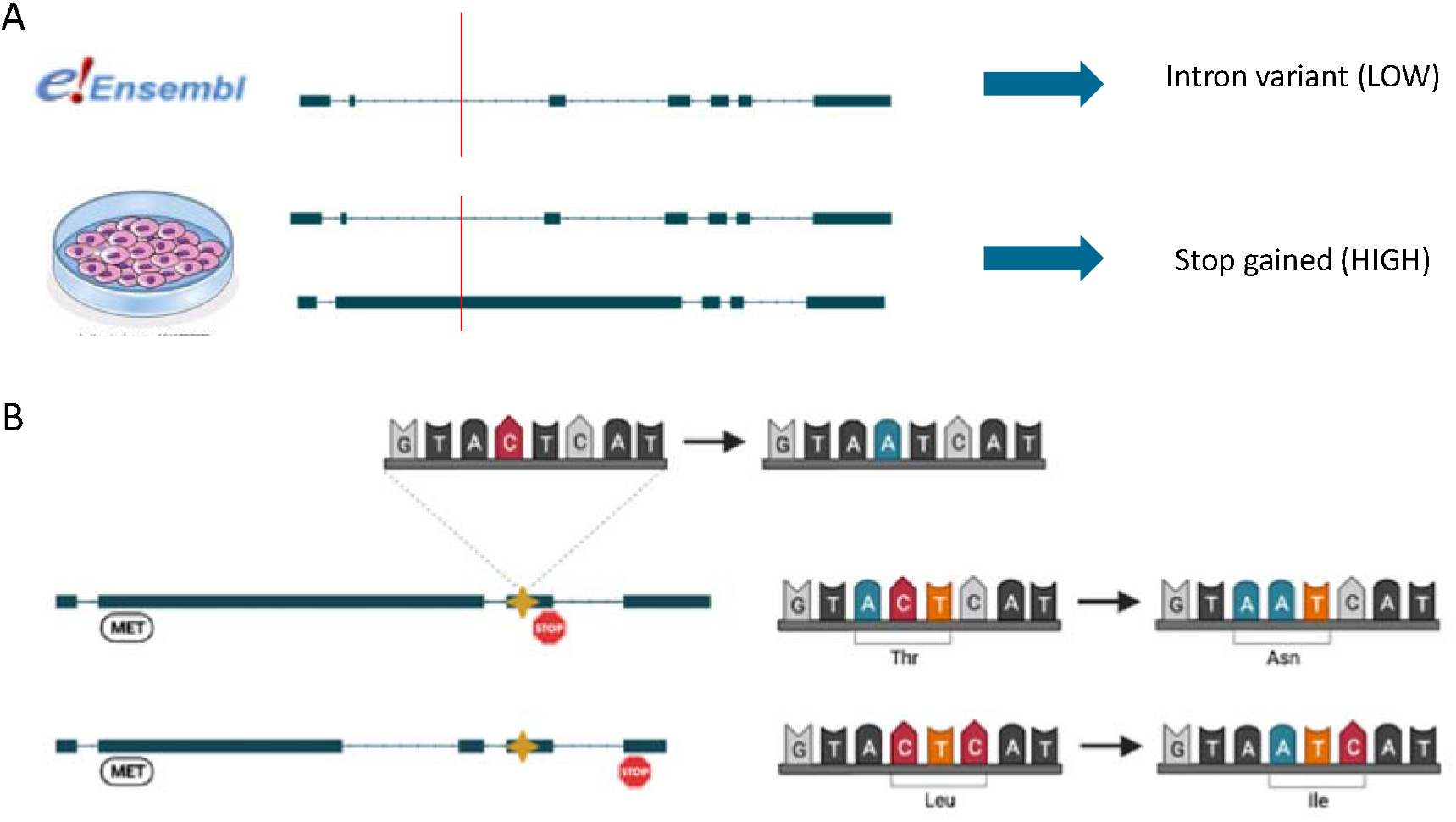
Premise for the creation of SUsPECT. A) Some pathogenic variants may be missed without actual transcript isoform information from a relevant sample. A variant in a particular genomic position may be incorrectly predicted to be non-deleterious. B) A variant at the same genomic position may cause a different missense variant in different transcript structures due to varying open reading frames per transcript.

The prediction of variant pathogenicity is an active area of development and, for ease of use, many tool creators release pre-computed sets of scores generated using reference transcript sets (27). This information is routinely used when evaluating variants against reference transcripts, but is not available when using novel transcript sets necessitating manual evaluation of the effects of variants on alternative proteoforms. One of the most commonly used variant annotators, Ensembl VEP, predicts molecular consequence for custom transcripts in standard formats, but the lack of pathogenicity predictions for missense variants in those transcripts limits interpretation. Considering the well-established importance of missense variants on a variety of diseases (28–30), this presents a hurdle in the re-annotation of variants with a custom transcriptome data.

The pipeline presented here, SUsPECT (Solving Unsolved Patient Exomes/gEnomes using Custom Transcriptomes), is designed to leverage cell/tissue-specific alternative splicing patterns to re-annotate variants and provide missense variant pathogenicity scores necessary for downstream variant prioritization. This pipeline was designed to be generalizable to any type of rare disease variant set paired with a relevant (long-read) transcriptome. For example, a researcher interested in annotating variants in a patient with a rare intellectual disability could consider using this tool along with a brain transcriptome dataset. We demonstrate the usefulness of this tool by reannotating ClinVar variants with a newly generated immune-related long-read RNA-sequencing dataset.

## Material and Methods

### Severity classification

SUsPECT classifies variants according to their expected impact and their molecular consequence. Impact scores used by SUsPECT are based on the predicted molecular consequence groupings in Ensembl VEP (Figure 2A) with higher numbers corresponding to more severe consequences: zero being equivalent to “modifier”, one to “low” severity, two to “moderate” severity, and four to “high” severity. SUsPECT uses Polyphen-2 predictions to distinguish between (likely) benign (score: 2) and (likely) deleterious (score: 3) missense variants.

**Figure 2:**
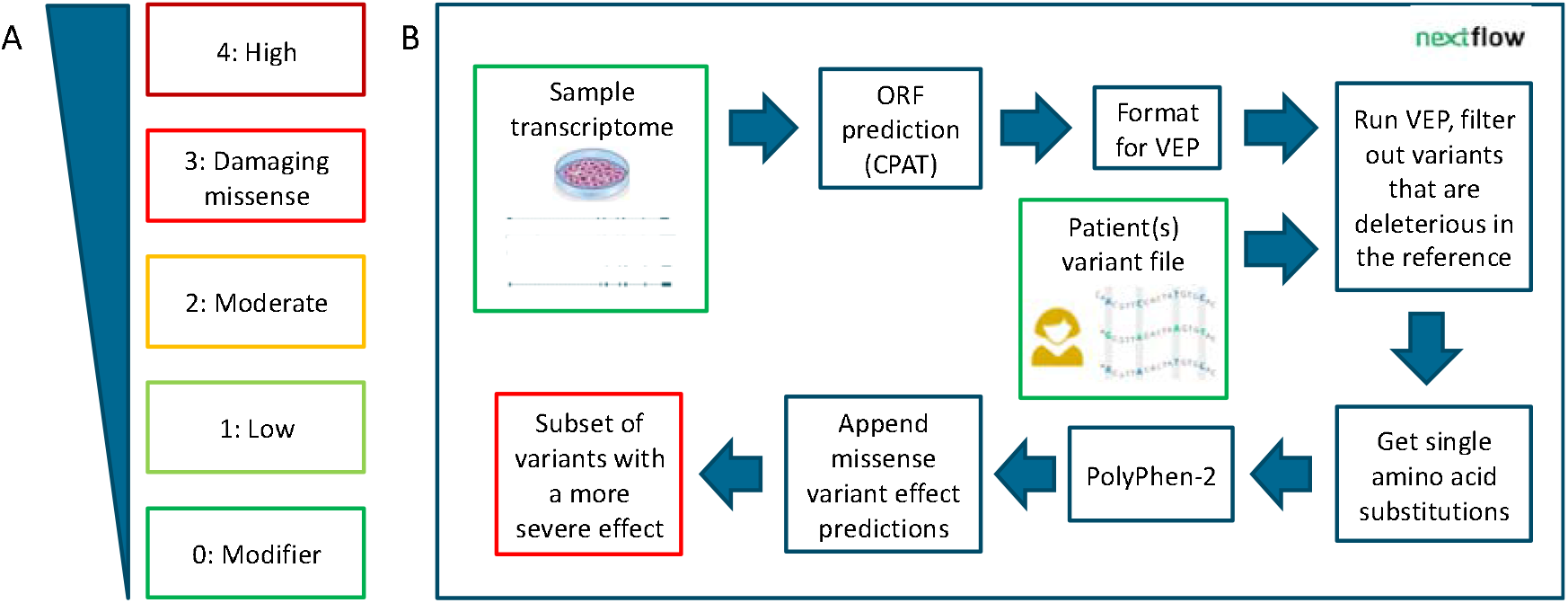
Reannotation with SUsPECT. A) Defining “more severe”. The five categories of severity are modifier, low, moderate, damaging missense and high. We consider levels 3 and 4 to be deleterious, and thus potentially pathogenic. B) The schematic of the pipeline.

### Additional filters for output variant list

SUSPeCT initial output is a list of variants with higher severity scores based on the custom transcriptome annotation compared to the reference annotation. The variants that remain in the final list of “increasing severity” are filtered to retain only variants that are potentially interesting for establishing a disease diagnosis. Thus, the pipeline removes variants that are already considered as (likely) pathogenic based on the reference annotation., i.e. variants that have original Ensembl VEP scores of 3 or 4. An additional criterion was applied for missense variants. Missense variants for which the same amino acid substitution found in the custom and reference annotation are also removed. To reduce computational time further, missense variant alleles in novel sequences that are common (AF > 0.01) are removed. These filters are integrated in SUsPECT. For the use case described in this manuscript, missense variants present in the custom annotation that are predicted by PolyPhen-2 to be “benign” in both custom and reference annotation are removed. In our ClinVar example, we define “immune-related” variants as those variants that contain the string “immun” somewhere in the clinical description.

### Software details

A pipeline was built to streamline the process of variant prioritization using custom transcript annotation. The pipeline is written in Nextflow (31), using Ensembl VEP as the variant annotator. Each step of the pipeline runs Singularity/Docker containers pulled automatically from Docker Hub. The input of the pipeline is the sample-specific/non-reference long-read transcriptome in GTF format, variants in a VCF file, and a FASTA file of the genome sequence. It is designed for use with output from TALON (32).

First, the GTF file is converted to BED format with AGAT v0.9.0(33). ORFs for any novel sequences are predicted based on the BED annotation and FASTA genome reference using CPAT v3.0.4. CPAT output is converted to BED format with the biopj python package and filtered for a coding probability of at least 0.364, which is the recommended human cutoff by the authors of CPAT(19). Conversion from CPAT CDS to protein FASTA is performed with EMBOSS transeq v6.5.7. This ORF BED file is combined with the BED file of transcripts to make a complete BED12 file with ORF/transcript information. Then, we convert this BED12 file to GTF with UCSC’s bedToGenePred and genePredToGtf. The resulting GTF file is used for a preliminary annotation of the variants with Ensembl VEP to fetch variants predicted as missense in the custom transcript sequences. Next, variant filtering was performed as outlined in the previous section with filter_vep utility distributed with Ensembl VEP as well as bedtools v2.30.0. The pathogenicity predictions are reformatted and one final run of Ensembl VEP (with the custom plugin enabled) integrates the pathogenicity predictions to the VCF. The output is the -annotated VCF, as well as a VCF with the subset of variants predicted to have higher severity.

### Ex vivo peripheral blood mononuclear cell (PBMC) experiments

Venous blood was drawn from a healthy control_(34)_ and collected in 10mL EDTA tubes. Isolation of peripheral blood mononuclear cells (PBMCs) was conducted as described elsewhere(35). In brief, PBMCs were obtained from blood by differential density centrifugation over Ficoll gradient (Cytiva, Ficoll-Paque Plus, Sigma-Aldrich) after 1:1 dilution in PBS. Cells were washed twice in saline and re-suspended in cell culture medium (Roswell Park Memorial Institute (RPMI) 1640, Gibco) supplemented with gentamicin, 50 mg/mL, 2 MM L-glutamine, and 1 mM pyruvate. Cells were counted using a particle counter (Beckmann Coulter, Woerden, The Netherlands) after which, the concentration was adjusted to 5 × 10^6^/mL. *Ex vivo* PBMC stimulations were performed with 5×10^5^ cells/well in round-bottom 96-well plates (Greiner Bio-One, Kremsmünster, Austria) for 24 hours at 37°C and 5% carbon dioxide. Cells were treated with lipopolysaccharide (*E. Coli* LPS, 10 ng/mL), *Staphylococcus aureus* (ATCC25923 heat-killed, 1×10^6^/mL), TLR3 ligand Poly I:C (10 µg/mL), *Candida albicans* yeast (UC820 heat-killed, 1×10^6^/mL), or left untreated in regular RPMI medium as normal control. After the incubation period of 24h and centrifugation, supernatants were collected and stored in 350uL RNeasy Lysis Buffer (Qiagen, RNeasy Mini Kit, Cat nr. 74104) at −80°C until further processing.

### RNA isolation and library preparation

RNA was isolated from the samples using the RNeasy RNA isolation kit (Qiagen) according to the protocol supplied by the manufacturer. The RNA integrity of the isolated RNA was examined using the TapeStation HS D1000 (Agilent), and was found to be ≥7.5 for all samples. Accurate determination of the RNA concentration was performed using the Qubit (ThermoFisher). Libraries were generated using the Iso-Seq-Express-Template-Preparation protocol according to the manufacturer’s recommendations (PacBio, Menlo Parc, CA, USA). We followed the recommendation for 2-2.5kb libraries, using the 2.0 binding kit, on-plate loading concentrations of final IsoSeq libraries was 90 pM (*C. albicans, S. aureus*, PolyIC, RPMI) and 100 pM (LPS) respectively. We used a 30h movie time for sequencing. The five samples were analyzed using the isoseq3 v3.4.0 pipeline. Each sample underwent the same analysis procedure. First CCS1 v6.3.0 was run with min accuracy set to 0.9. Isoseq lima v2.5.0 was run in isoseq mode as recommended. Isoseq refine was run with ‘--require-polya’. The output of isoseq refine was used as input for TranscriptClean v2.0.3. TranscriptClean was run with ‘--primaryOnly’ and ‘--canonOnly’ to only map unique reads and remove artifactual non-canonical junctions of each of the samples. The full TALON pipeline was then run with all five samples together using GRCh38 (https://www.encodeproject.org/files/GRCh38_no_alt_analysis_set_GCA_000001405.15/@@download/GRCh38_no_alt_analysis_set_GCA_000001405.15.fasta.gz). Assignment of reads to transcripts was only allowed with at least 95% coverage and accuracy. A minimum of 5 reads was required to allow isoforms to be kept in the final transcript set (default of talon_filter_transcripts). Ensembl/GENCODE annotation (v39) was used by TALON to determine novelty of transcripts in the sample.

## Results

### Analysis pipeline overview

We developed SUsPECT to re-annotate variants using custom transcriptomes. This pipeline returns a VCF file with alternative variant annotations for downstream evaluation and prioritization. SUsPECT is based on Ensembl VEP and additionally predicts pathogenicity for missense variants different from the user-provided RNA sequencing dataset. A schematic overview of the pipeline is presented in Figure 2B. The main steps in the pipeline are:

- Validate pipeline input, including 1) an assembled (long-read) transcriptome in GTF format with novel transcripts. A long-read transcriptome assembly tool such as TALON will output a suitable file. 2) A VCF containing patient(s) variants.
- ORF prediction is performed on the transcripts that do not match any in the human reference transcriptome.
- Ensembl VEP adds predicted molecular consequence annotations based on your transcripts/ORFs. Variants considered as missense in the user-provided transcriptome are reformatted and submitted to Polyphen-2.
- Polyphen-2 calculates pathogenicity scores and provides predictions. These are reformatted and incorporated into the final VCF annotation file.
- A sub-list of variants that have a more severe molecular consequence in the input transcriptome are provided in tabular format.

### A long-read sequencing transcriptome of stimulated PBMCs

We have generated long-read sequencing data on atypical, *i*.*e. in vitro* stimulated samples - provoking a strong expression response, to illustrate the use of the pipeline. We chose this dataset to exemplify less-studied tissues/conditions because novel transcripts are more numerous in these samples and SUsPECT is most likely to yield interesting results when the input transcriptome has many novel transcripts. Our custom transcriptome is based on long-read transcript sequences related to host-pathogen interactions and is derived from human PBMCs exposed to four different classes of pathogens. We combined the transcript structures of all four pathogenic conditions and control samples for the reannotation. We identified a total of 80,297 unique transcripts, 37,434 of which were not present in the Ensembl/GENCODE reference transcriptome. Relative abundances of novel transcripts were lower than of reference transcripts (Supp figure 1). The custom transcriptomes resulted in prediction of 34,565 unique novel ORFs passing CPAT’s coding capacity threshold. The majority of transcripts had at least one ORF predicted (Supp figure 2).

### Reannotation of ClinVar variants

Variants that fall in the novel transcripts may result in a more severe molecular consequence, but the functional and ultimately clinical implications remain unclear. We therefore focused on re-annotating ClinVar variants to demonstrate that SUsPECT can suggest new candidate pathogenic variants associated with clinical outcomes. ClinVar contains variants with clinical significance curated by different authorative sources. We hypothesized that ClinVar variant that were annotated as pathogenic and not predicted to to be deleterious with the reference annotation, but predicted deleterious with a (relevant) sample transcriptome, would support the utility of this pipeline.

We tested SUsPECT on a recent ClinVar (36) release (April 2022), excluding all variants that were annotated in ClinVar to be (probably) benign. We compared the predicted severity of the 776,866 variants using our custom transcript annotation versus the reference. After applying filters as described in the Methods section, 1,867 candidate variants remained. Of these variants, 145 were associated with monogenic immune-related disorders, which is significantly more than expected by chance (odds ratio=5.46, p=1.51×10^−55^). This could indicate that annotation with an immune-relevant transcriptome is better suited for the identification of variants with an impact on immune function than annotating with a reference transcriptome. The strongest argument for the utility of this pipeline can be made with variants that are curated in ClinVar to be pathogenic rather than those of uncertain significance (VUS). After excluding variants of unknown significance (VUS) from the full candidates list, there are 90 variants remaining (5 immune-related). These 90 variants had an enrichment of severity level 4 events (Supp figure 3).

Five immune-related variants curated in ClinVar to be pathogenic were reannotated from a low severity molecular consequence in the Ensembl/GENCODE transcript set to a moderate or high severity in our transcriptome (Table 1). Two were missense variants in the custom annotation and three were start-loss/stop-gain. We visualized the variants in the context of the transcript structures/ORFs on the UCSC genome browser. Two examples can be seen in Figure 3. The variant in *IFNGR1* (dbSNP identifier rs1236009877) is associated with IFNGR1 deficiency. It is curated by a single submitter in ClinVar as ‘likely pathogenic’ using clinical testing. Annotation of the variant with reference transcripts results in a low severity (intronic variant) result, but results in a stop-gain variant (high severity) when annotating with our transcriptome. Our custom transcriptome contained multiple novel transcripts with a retained intron at the site of the variant, but only 1 of these transcripts had a predicted ORF in this intron. The particular transcript affected by this stop gained variant was found in all samples sequenced with minimum 3 and up to 10 supporting reads, indicating that it is unlikely an artifact. The predicted ORF extended 30 base pairs into the retained intron in the region of this variant. It was the most probable ORF for that transcript with a coding probability by CPAT of 0.934.

**Table 1:** Five ClinVar pathogenic immune-related variants were reannotated from low severity in hg38 to high severity in the custom transcriptome.

| dbSNP | Location hg38 | Allele | Gene | Consequence reference | Consequence custom | ClinVar condition |
| --- | --- | --- | --- | --- | --- | --- |
| rs80358236 | 1:172665641 | C | FASLG | In-frame deletion | Start lost & in-frame deletion | Autoimmune lymphoproliferative syndrome |
| rs1573262398 | 2:97724319 | T | ZAP70 | Benign missense | Missense (unknown) | Combined T and B cell immunodeficiency |
| rs113994173 | 2:97733464 | A | ZAP70 | Intron | Missense (unknown) | Combined immunodeficiency due to ZAP70 deficiency |
| rs387906763 | 2:190999647 | G | STAT1 | Benign missense | Start lost | Immunodeficiency 31C |
| rs1236009877 | 6:137203727 | A | IFNGR1 | Intron | Stop gained | Immunodeficiency 27A |

**Figure 3:**
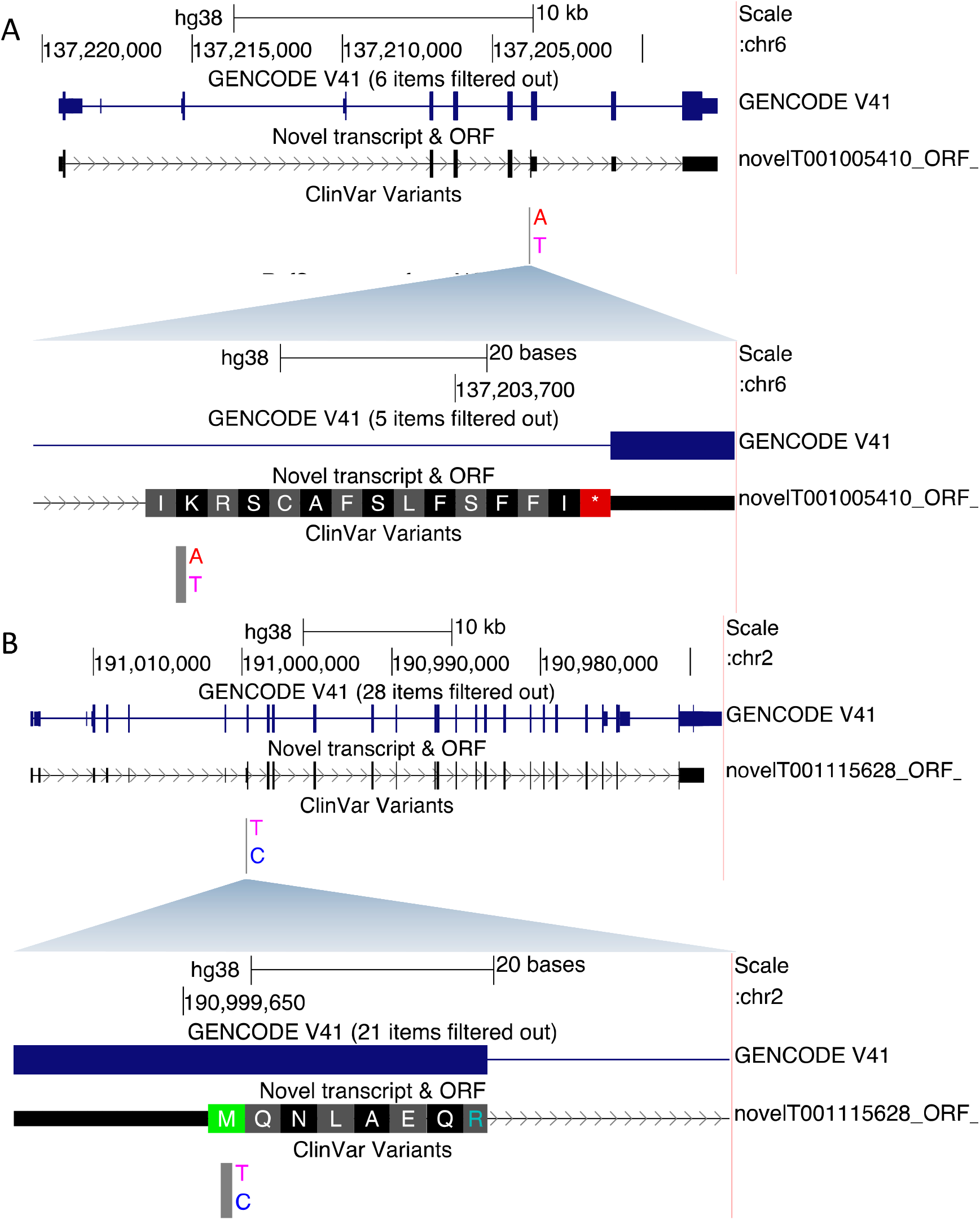
Two examples of ClinVar pathogenic variants being re-annotated. Both variants were considered low severity variants when using hg38 reference transcriptome to annotate. A) *IFNGR1* whole view and close-up of region around the variant. Variant causes a stop-gain effect (K>*) in the custom transcript novelT001005410. B) *STAT1* whole view and close-up of region around variant. Variant causes a start loss (M>T) in the custom transcript novelT001115628.

In addition, the variant in *STAT1* (dbSNP identifier rs387906763) was pathogenic according to the LitVar (37) literature mining tool and a clinical testing submission. It is a missense variant (Tgc/Cgc) in the reference annotation that is predicted by PolyPhen-2 to be benign. However, in one novel transcript it causes an M/T substitution, leading to loss of translation start site. Further inspection revealed that the transcript affected by the start-loss was expressed in *C. albicans, S. aureus* and PolyIC stimulated conditions by up to 6 supporting reads, but 0 in the control condition. STAT1 is previously described to be involved in the immune disease (chronic mucocutaneous candidiasis) linked to this variant by weakened response to *C. albicans* (38), which is a condition where this novel transcript was expressed. The ORF affected was the most probable ORF for that transcript and had a coding probability of almost 1 by CPAT.

## Discussion

The human transcriptome is more complex than the current reference annotation would suggest. Variants in non-reference transcripts may aid in explaining missed genetic diagnoses, especially when disease-specific transcripts are used. SUsPECT puts genetic variants in the context of transcript isoform expression and can contribute to an increase in diagnostic yield. We used ClinVar pathogenicity assertions to demonstrate the potential of this methodology to re-annotate variants that may have previously been overlooked due to insufficient transcript isoform information. We have shown that annotating missense variants in the light of the expressed isoforms can change their predicted effect from benign to pathogenic. The enrichment of immune-related variants after reannotation suggests there is biological significance to these findings.

Considering the clinical applications of this pipeline, it is important to underline that variant causality is not an output of this pipeline. The pipeline simply brings new candidates forward for further interpretation; the user may choose to cross-reference the clinical phenotypes of the patients with the functions of the genes that the patients’ variants are found to disrupt. In our use case, ClinVar variants were used as they already have widely accepted annotations. However, 40% of ClinVar is made up of variants of unknown significance (VUS), some of which are suspected to have some impact on clinical phenotype. Many of these variants changed annotation from benign to deleterious in our reannotation. As more people use sample-specific transcriptomes to annotate variant sets, an increasing number of VUS may be classified as benign or deleterious.

We observed that many increased severity variants were missense, which may have to do with the numerous new ORFs. Multiple ORFs passing CPAT’s ‘human threshold’ were often predicted per novel sequence; for our 37,434 novel sequences we predicted 34,565 novel ORFs. Some proteogenomics tools choose the ‘best’ ORF per sequence, but we have chosen to keep all that passed the probability threshold. We do not filter out non-coding genes when predicting ORFs, opting instead for minimal filtering to provide all information to the end user. Missense results implicitly depend on the confidence of the ORF predictions that are produced by CPAT. New deleterious missense variants may not be relevant if the protein in question is not produced. Coding ability of novel transcript isoforms is an area of active research (39–41) and new techniques to identify credible ORFs may be added to the pipeline as they become available. In the meantime, it may be prudent to validate interesting candidates using targeted proteomics techniques before establishing a genetic diagnosis.

## Data Availability

SUsPECT is open source and freely available for download on Github (https://github.com/cmbi/SUsPECT)

Raw PacBio sequencing data and transcriptome is available on EGA under accession number ##.

## Acknowledgement

We would like to thank Simon V. van Reijmersdal for his contribution to the library preparation.

