## Supplemental figures for "SUsPECT: A pipeline for variant effect prediction based on custom long-read transcriptomes for improved clinical variant annotation"

Supplementary Data

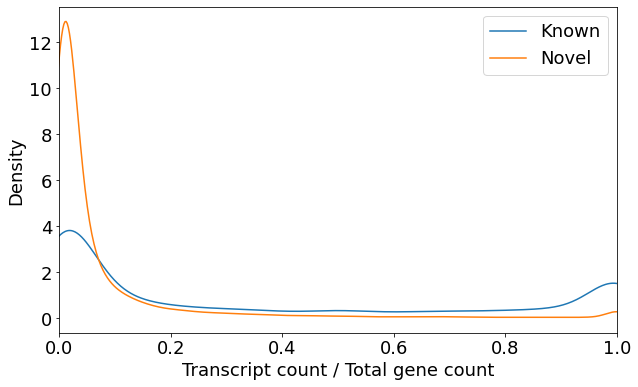

Supplementary figure 1: Relative abundance of novel transcripts relative to known transcripts. Transcript counts were summed for all 5 conditions per transcript.

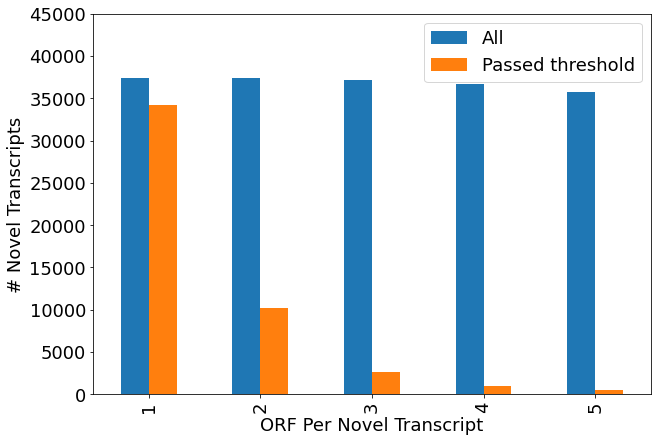

Supplementary figure 2: Number of predicted ORFs passing CPAT’s human coding threshold. ORFs were predicted per novel transcript and sorted by most to least likely to be coding (1 being most likely). All predicted ORFs in blue, those that passed the human coding threshold in orange.

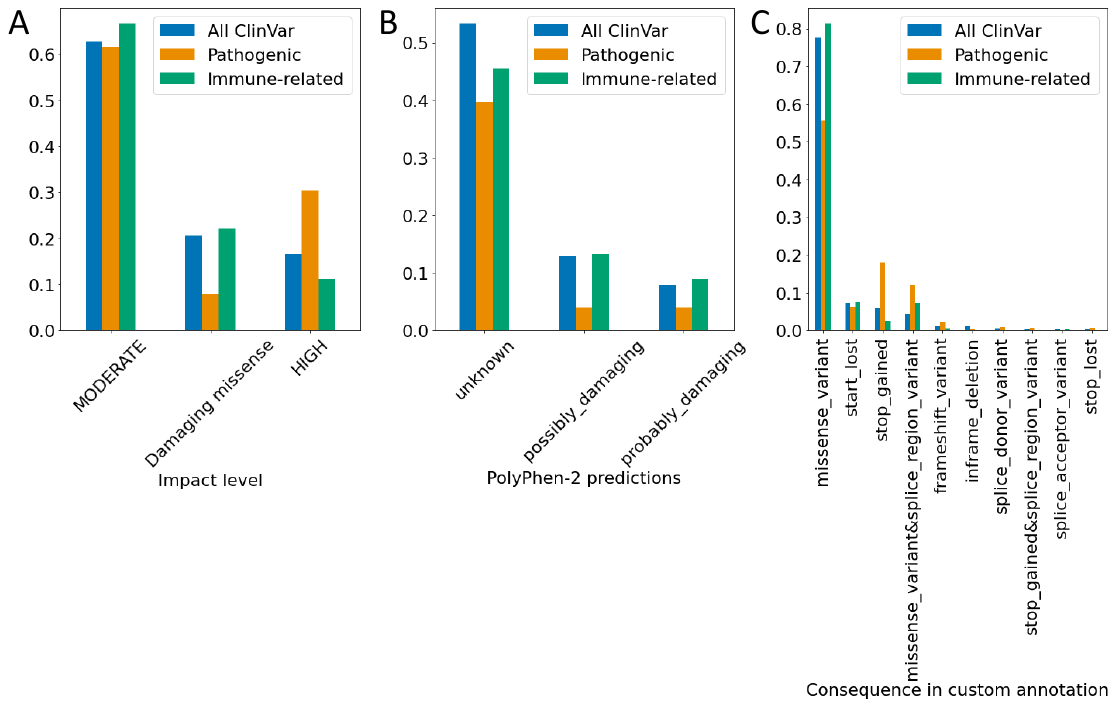

Supplementary figure 3: Comparing subsets of variants that changed annotation from benign to deleterious. “All ClinVar” corresponds to all reannotated clinvar variants including VUS and all clinical phenotypes that were reannotated from benign to deleterious with our transcriptome (N=1867). “Pathogenic” is a subset of all reannotated variants that excludes VUS variants (N=90). “Immune-related” is a subset of all reannotated variants that includes only immune-related clinical phenotypes (N=145). A) Impact level of variants after reannotation. The impact level shown is associated with annotation in the custom transcriptome. B) PolyPhen-2 predictions of variants after reannotation. C) Specific molecular effects of variants after reannotation.
